# Genotyping of structural variation using PacBio high-fidelity sequencing

**DOI:** 10.1101/2021.10.28.466362

**Authors:** Zhiliang Zhang, Jijin Zhang, Lipeng Kang, Xuebing Qiu, Beirui Niu, Aoyue Bi, Xuebo Zhao, Daxing Xu, Jing Wang, Changbin Yin, Xiangdong Fu, Fei Lu

## Abstract

**Background:** Structural variations (SVs) pervade the genome and contribute substantially to the phenotypic diversity of species. However, most SVs were ineffectively assayed because of the complexity of plant genomes and the limitations of sequencing technologies. Recent advancement of third-generation sequencing technologies, particularly the PacBio high-fidelity (HiFi) sequencing, which generates both long and highly accurate reads, offers an unprecedented opportunity to characterize SVs and reveal their functionality. Since HiFi sequencing is new, it is crucial to evaluate HiFi reads in SV detection before applying the technology at scale.

**Results:** We sequenced wheat genomes using HiFi, then conducted a comprehensive evaluation of SV detection using mainstream long-read aligners and SV callers. The results showed the accuracy of SV discovery depends more on aligners rather than callers. For aligners, pbmm2 and NGMLR provided the most accurate results while detecting deletion and insertion, respectively. Likewise, cuteSV and SVIM achieved the best performance across all SV callers. We demonstrated that the combination of the aligners and callers mentioned above is optimal for SV detection. Furthermore, we evaluated the impact of sequencing depth on the accuracy of SV detection. The results showed that low-coverage HiFi sequencing is capable of generating high-quality SV genotyping.

**Conclusions:** This study provides a robust benchmark of SV discovery with HiFi reads, showing the remarkable potential of long-read sequencing to investigate structural variations in plant genomes. The high accuracy SV discovery from low-coverage HiFi sequencing indicates that skim HiFi sequencing is an ideal approach to study structural variations at the population level.

## Background

Structural variations (SVs) and single nucleotide polymorphisms (SNPs) are two ends of the genetic variation spectrum. On the contrary to the simplicity of SNPs, SVs exhibit a much higher level of complexity—insertion, deletion, duplication, inversion, and translation, varying in size from ~50 bp^1^ to hundreds of megabases (Mb), constitute a highly diverse set of SVs in the genome^234^. While SVs being considered as a major source of casual variation in crop traits, such as flowering time in maize (e.g., *Vgt1*^5^, *ZmCCT*^6,7^), the grain yield in rice (e.g., *GW5*^8^ and *GL7*^9^), the solid-stemmed architecture in wheat (e.g., *TdDof*^10^), and the smoky volatile locus in tomato (e.g., *NSGT1*^11^ and *NSGT2*^12^), the detection and genotyping of SVs remains to be one of the greatest challenges in genomic studies^13^.

Aside from the structural complexity of the genome, technological limitations are also restricting SV detection and genotyping^14,15^. Even though the cost of Next-generation high-throughput sequencing (HTS) remains relatively low, it is constrained by the short read length, causing the insufficient power to detect the large SVs. Long-read sequencing, such as PacBio CLR and Oxford Nanopore, has the advantage of scanning large SV, but their high base error rate (~ 8 - 20%) put forward higher demands for long reads aligner and SV caller, which restricting strongly the wide application owing to imprecise breakpoint and inaccurate SV sequence^16,17^. Encouragingly, the PacBio CCS method generates high accurate long HiFi reads (>10 Kb) and seems to strike the perfect balance between reads accuracy and length, further improving SV detection^18^. Meanwhile, the corresponding algorithms, such as aligners (pbmm2, NGMLR^19^, or Minimap2^20^) and callers (pbsv, cuteSV^21^, SVIM^22^, and Sniffles^19^), were developed for long-scale SV identification, though almost all the SV algorithm were generally designed for the diploid human genome. The progress of sequencing technology will bring the great reform in SV detection, to which we say: 1) how to establish robust benchmark tools for SV discovery based on HiFi reads; 2) how to achieve the higher performance to deeply mine the SV in the plant genome; 3) how to optimize parameter on SV calling to deal with the relatively high cost at the population level.

Here, we provide the first workflow with general applicability to evaluate SV detection using current long-read aligners and SV callers based on HiFi reads. The analysis establishes not only a robust guideline for SV detection with HiFi reads in the plant genome, but also the parameter optimization for low-coverage data mining. Predictably, this study will facilitate the large-scale application of PacBio HiFi sequencing technology at the population level.

## Results

### Schematic workflow of evaluating SV detection algorithm

We evaluated the performance of existing long-read-based SV callers and alignment programs against the ground truth set identified by integrating multiple genomic data in the following approach (Fig. 1). In step1 (“SV calling”), SV sets were obtained using pbsv, cuteSV, SVIM, and Sniffles after pbmm2, NGMLR, or Minimap2 alignment. Because pbsv cannot recognize the Minimap2 alignment format, we finally identified a total of 11 SV sets with combinations of the aligners and SV callers. In step2 (“Truth set construction”), due to the lack of available ground truth SV set in the wheat genome, several samples were deep re-sequenced. For each candidate SV, the discordant alignment features and depth features were characterized by the methods of read depth (RD), read pair (RP), and split read (SR) from short-read sequencing data (see the “Methods” section for details). Therefore, the truth SV set was formed based on the data integration of multiple sequence technologies, which is close to comprehensively characterizing SVs, although it was devoid of undiscovered SVs. In step3 (“Precision-recall comparison”), we were able to test the performance of 11 SV sets by estimating precision, recall, and F-measure using the truth SV set. Overall, we provide a general workflow for comprehensive evaluation of long-read aligners and SV callers, hoping to build a robust benchmark for SV detection.

**Fig. 1.**
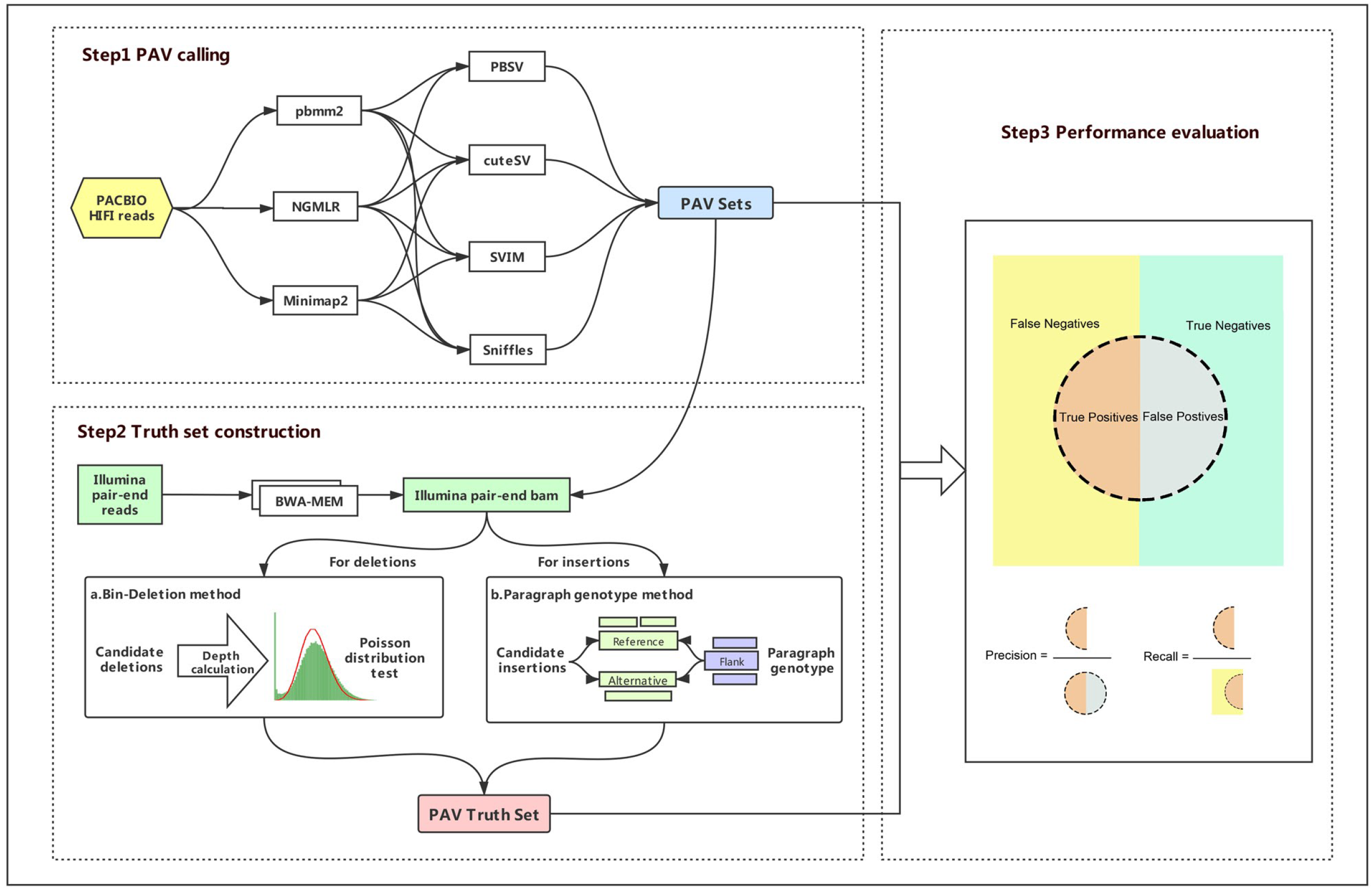
Schematic workflow of the comprehensive evaluation of long-read aligners and SV callers. **Step 1 SV calling** SV sets were obtained from the 4 callers and 3 aligners combination of pairs based on PacBio HiFi reads. **Step 2 Truth set construction** Truth SV set was formed based on the data integration of multiple sequence technologies (see the “Methods”). **Step 3 Performance evaluation** Comprehensive evaluation of aligners and callers for Pacbio HiFi long-reads.

### Sequence-resolved candidate sets of structural variation (SV)

To facilitate the study of genome-wide identification of SVs in different wheat accessions, we tested three ploidy levels (AABBDD, AABB, DD genome) from *Triticum/Aegilops* using PacBio Circular Consensus Sequencing (CCS) mode, generating highly accurate HiFi reads with an average length of 13.0 kb, 17.2 kb, 12.9 kb, respectively (Additional file 2: Tables S1, Additional file 1: Figure S1). By applying our previously designed pipeline for cross-ploidy genetic variation discovery, we identified 11 sequence-resolved candidate SV sets per sample from the combination of the aligners and SV callers^23,24^. In general, all SV callers were similar with respect to the number of SVs after NGMLR or pbmm2 alignment, but the SVs count by Minimap2 is higher than the other two aligners (Fig. 2a, Additional file 1: Figure S2a, Additional file 2: Tables S2). As expected, the SV size distribution showed decreasing frequency with increasing length and was deeply affected both by aligner and caller (Fig. 2b). For deletion, more smaller (50-100bp) events could be detected by aligner Minimap2. Also, NGMLR could detect the little large events(~300bp) for insertion.

**Fig. 2.**
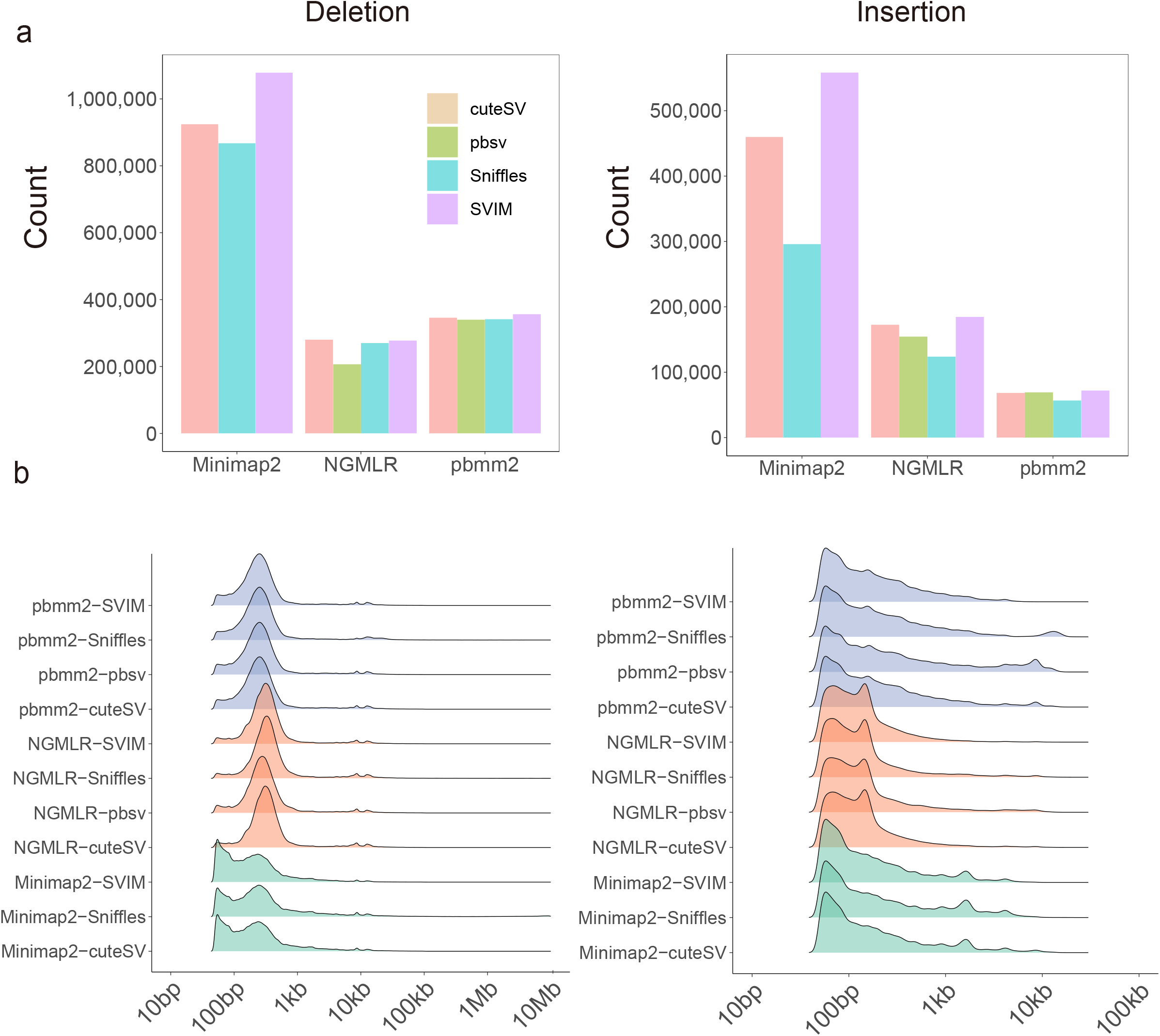
Summary of the sequence-resolved candidate SV sets. The number (**a**) and size distribution (**b**) of each SV type, deletion and insertion, from 11 sequence-resolved candidate SV sets.

The occupancy of computing resources, an important factor considered by users including run time and average memory usage, was then examined using 20 threads in the hexaploid genome (Additional file 1: Figure S2, Additional file 2: Tables S3). For run time, aligner pbmm2 and Minimap2 generally processed the same datasets (~30Gb) with 5-7 times less runtime relative to NGMLR (380.4 & 562.8 mins vs. 2691 mins), and callers pbsv took a long time for SV detection than the other three callers. For average memory usage, aligners occupied relatively high memory (32-50 G). In addition, callers cuteSV, Sniffles, and SVIM used a similar memory (≤□6□G), and pbsv required a little more memory (~16 G). Overall, the impact on computing resources was mainly concentrated in the aligner, not the SV caller.

### The base-level SV truth set

Unlike human, little SV study is relatively available in the wheat genome. To evaluate the performance of the SV detection algorithm, the SV truth set was first constructed using data integration of multiple sequence technologies^25^. By deep re-sequencing (14~25 X) of these samples, we had the ability to validate the results of each 11 SV sets through utilizing the discordant alignment and depth features (Additional file 2: Tables S1). For deletion, we developed an efficient pipeline, Bin-deletion, by calculating the depth features for which deletions were discovered. Due to a large number gap in the wheat genome, we corrected depth and chose adjustDepth = 0 for the deletion truth set (see the “Methods” section for details). Given the success of genotyping tools of structural variations (SVs)^26,27,28^, we use paragraph^26^, an accurate genotyper for short-read sequencing data, to further validate the insertion dataset that had been mined by long-read HIFI data.

In addition, many packages for merging structural variants (SVs) among multi-VCF files have been released in recent years^29,30^. It is worth noting that previous work rarely incorporated the effect of a maximum allowed merging distance, which usually used 500 or 1000 bp distance, resulting in a decline in the number of SVs and imprecise breakpoint position^31,32,33^. Moreover, there are obvious distinctions among callers for the same candidate insertion. To avoid these issues and obtain a more accurate truth set, each truth set of the corresponding candidate SV set was independently constructed, respectively.

For deletion, all deletion truth sets obtained by the method of Bin-deletion were merged to form the deletion truth set. For insertion, due to the difference in callers, we then tested the position distance for the two adjacent records, which had a 2-bp difference in size for insertion sequence after merging multiple insertion truth files identified by paragraph. 2-bp distance in the left and right breakpoints, which was able to combine 95.83 % of the nearest insertion data within a 2-bp length difference, was permitted for merging or comparing any insertion files. Combined with the above results, the base-level SV truth set was formed and considered as a real dataset for further analysis (Additional file 1: Figure S3, Additional file 2: Tables S4).

### The impact of aligners and callers on SV detection

No previous study to discuss the impact of aligner and caller on the accuracy of the SV set. To investigate this, we first constructed a total of 63 SV sets based on aligner, including 30 two-callers, 18 three-callers, and 4 four-callers SV sets obtained by the integration of caller pbsv, cuteSV, SVIM, and Sniffles after pbmm2, NGMLR and Minimap2 alignment, respectively, as well as 11 single-caller SV sets each from the caller and aligner combination of pairs. Furthermore, 37 SV sets based on the same caller, composed of 11 single-aligner, 20 two-aligners and 6 three-aligners, were obtained from the combination (intersection or union) of multiple SV sets.

According to the aligners, 63 SV sets, either deletion or insertion, were clearly divided into three groups (Fig. 3). There was an obvious contrast in precision among aligner-based methods, but not the 37 caller-based SV sets. Robust analysis of variance for 11 single-caller SV sets in different ploidy levels indicated that the result was a significant difference across aligner and caller (Table 1). The variance explained by the aligner was greater than the variance explained by the caller, especially the precision of SV sets. The results showed that both the precision and recall varied depending on aligner rather than caller, so 63 SV sets based on aligner should be recommended for in-depth analysis.

**Fig. 3.**
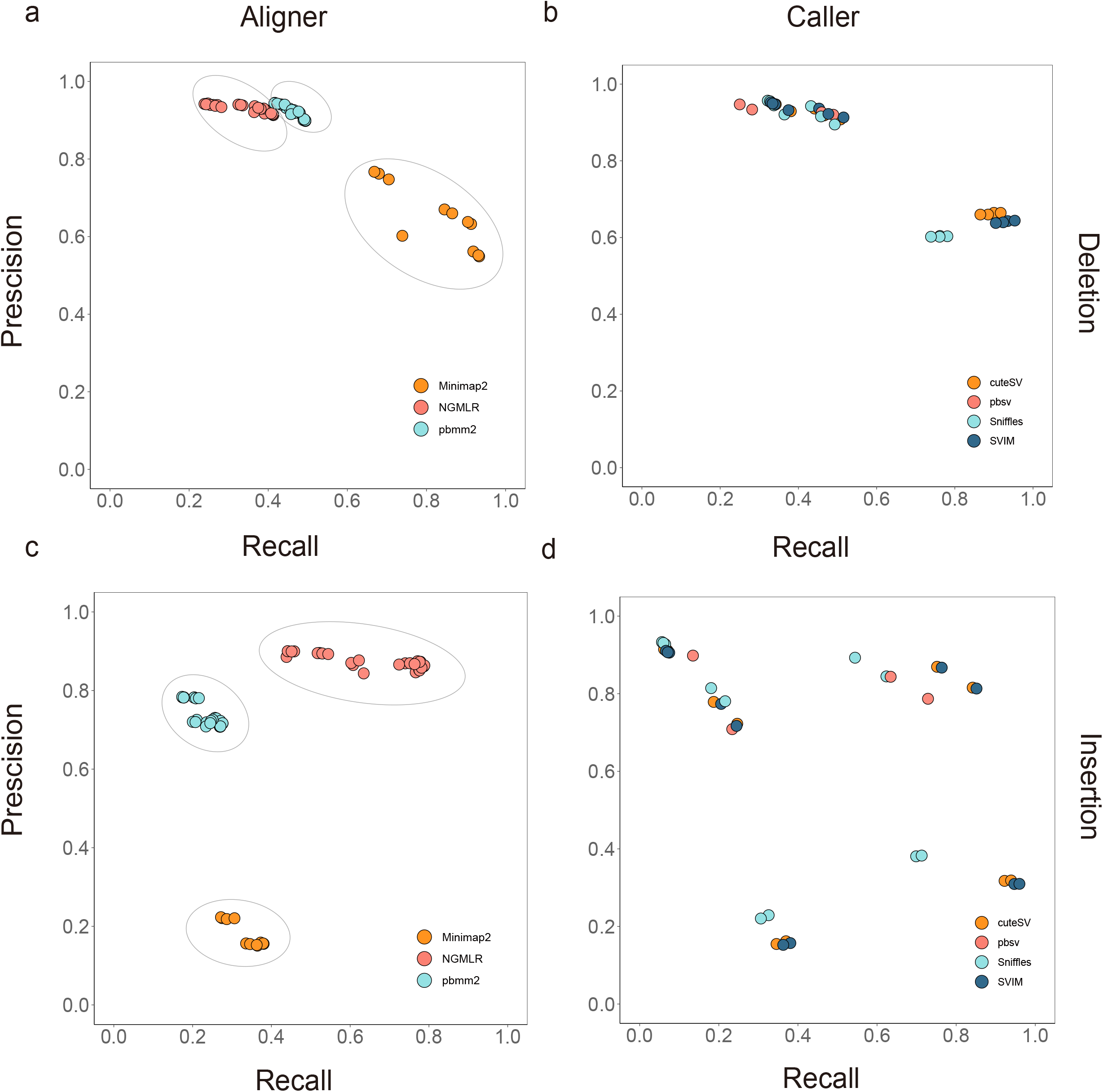
The impact of aligners and callers on SV detection. **a** (deletion) **and c** (insertion) Precision-recall plot of 63 SV sets obtained by a combination of multiple callers based on the same long-read aligner. **b** (deletion) **and d** (insertion) Precision-recall plot of 37 SV sets obtained by a combination of multiple aligners based on the same SV caller.

**Table 1.**
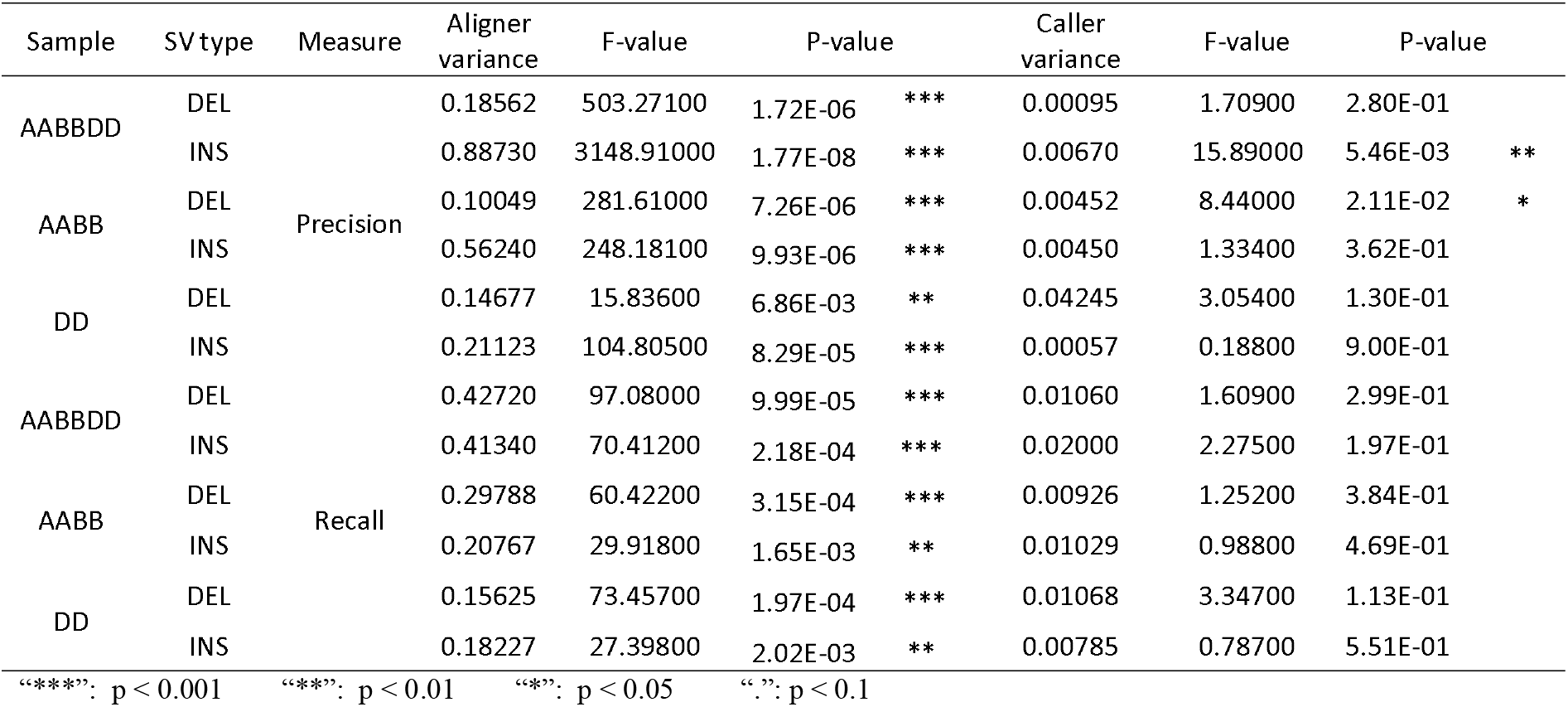
F-test between aligners and callers.

#### Overall performance of 63 SV detection algorithm

Establishing a standard method for SV detection posed big challenges for users in algorithm selection based on HiFi read. Recent research showed that a combination (intersection or union) of multiple SV callers could contribute to obtaining confidence or sensitivity results based on Illumina short-read^34,35^ and Oxford Nanopore long-read data^17^. Considering the advantage of highly accurate long HiFi reads, can this combination approach of SV callers improve the precision/recall? We further integrated a total of 63 SV sets based on aligner to evaluate the effect of single/combining SV call sets against the base-level SV truth set in hexaploidy (AABBDD) genome.

Minimap2 worked less well in terms of precision than the other aligners, either deletion or insertion, and aligner NGMLR or pbmm2 were emphatically discussed for further analysis (Fig. 4, Additional file 2: Tables S5-7).

1. Single SV detection algorithm: For deletion, the highest F-measure was obtained using cuteSV, SVIM or pbsv after pbmm2 alignment, and Sniffles was less powerful resulting in a lower precision (Fig. 4a, Additional file 1: Figure S4a S5a). For insertion, callers cuteSV or SVIM also achieved good performance, but NGMLR was more accurate than aligner pbmm2, which was a great deal of difference compared to deletion (Fig. 4b, Additional file 1: Figure S4d S5d).
2. Combining SV detection algorithm: As expected, the recall values of high-confidence (intersection) sets gradually decreased with the increase of combined SV sets (Fig. 4, Additional file 1: Figure S6a S7d, Additional file 2: Tables S6). However, given the similar precision values for deletion, insertion showed a general trend with the apparent addition in the values of precision compared with single caller SV set in three ploidy levels. On the contrary, high-sensitivity (union) sets, generated using two or more SV sets, could be capable of increasing recall with a bit of change of precision for both deletion and insertion (Fig. 4, Additional file 1: Figure S7a S7d, Additional file 2: Tables S7).

**Fig. 4.**
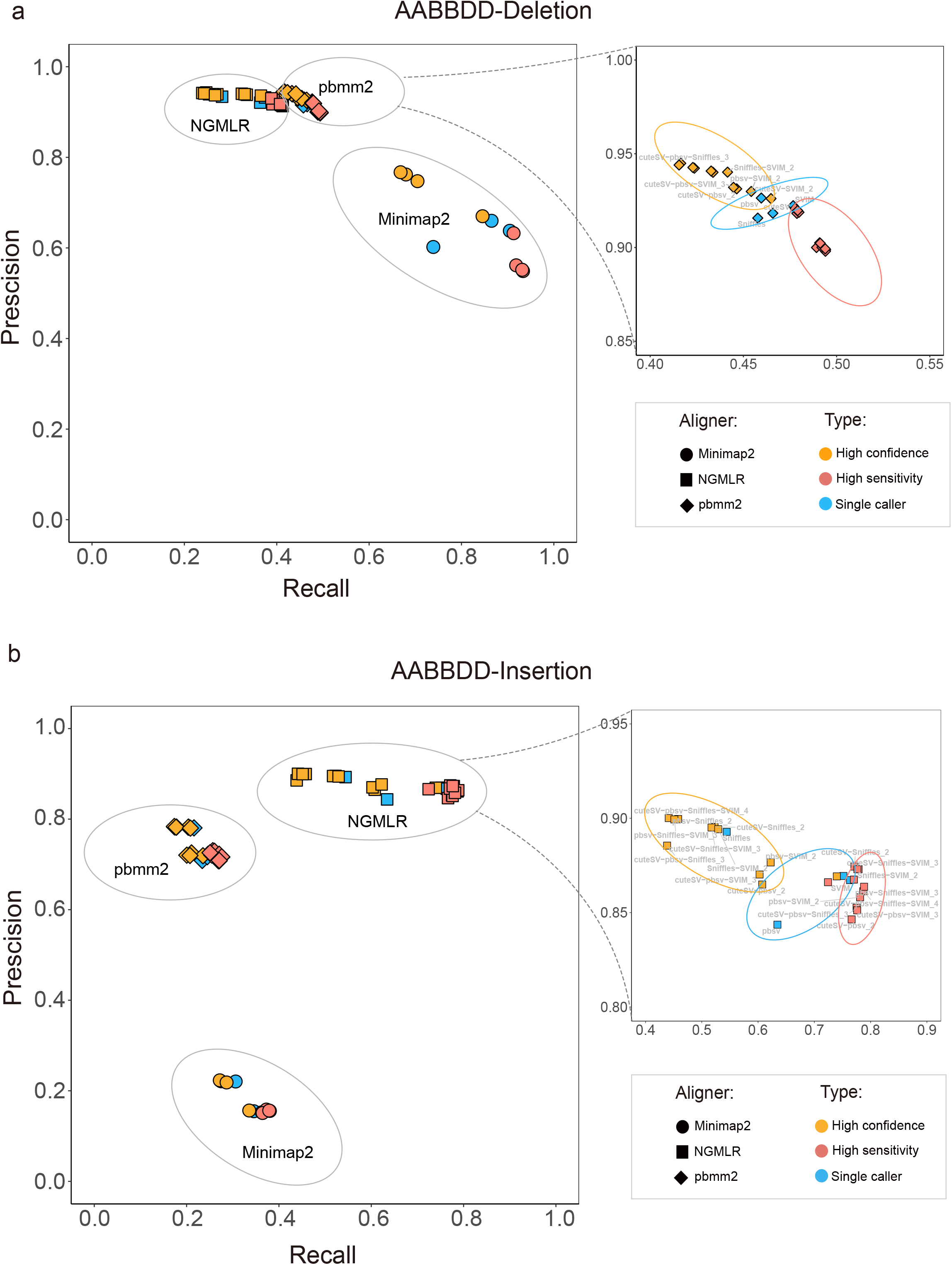
Comprehensive evaluation of 63 SV sets in hexaploid (AABBDD) genome. **a** (deletion) **and b** (insertion) Precision-recall graph of single/combining SV call sets against the base-level SV truth set. Aligners are represented by symbols, and multiple set sources are represented by colors as specified in the legend.

In summary, benchmark tools for SV detection could be recommended that caller cuteSV or SVIM after aligner pbmm2 (for deletion) or NGMLR (for insertion) achieved the optimum performance for the HiFi data in hexaploidy genome. High-confidence results could be obtained by combining multiple SV callers for insertion on precision, and could not be significantly improved in deletion calling. However, high-sensitivity results, both deletion or insertion identification, were significantly increased on recall values (Additional file 1: Figure S7a S7d, Additional file 2: Tables S7).

### Benchmark tools should be independent of ploidy level

Almost all the SV algorithms are designed to detect large-scale genomic variation for the diploid human genome^2^. However, the nature of the plant genome, distinct differences in ploidy variation^36^, present challenges to deeply mine the character of SV variation. To test the effect of ploidy levels on SV calling, we then analyzed the tetraploid (AABB) and diploid (DD) genome applying the above method (Additional file 1: Figure S8 S9). Like the hexaploid genome, precision-recall curves presented clear information that caller cuteSV or SVIM exhibited higher performances in calling SV and aligner pbmm2 or NGMLR achieved the best performances for the deletion or insertion data, respectively. Furthermore, more confident or sensitive results could be obtained by a combination of overlapping SV callers. These results demonstrated that the performance, including precision and recall, was entirely irrelevant to the ploidy level (Additional file 1: Figure S10).

### Impact of sequencing depth on precision

PacBio CCS method produces highly accurate (~99.8%) and long (>10 kb) reads, which greatly enhancing the ability of SV detection^18^. However, this approach is difficult to scale up for SV genomics studies at the population level, due to its relatively high cost and lower output of HiFi data, especially in the large and hexaploidy wheat genome. To lower these limitations and obtain confidence results, it is particular importance to evaluate the relation between sequencing depth and precision.

Given the relative higher depth in the diploid (DD) sample, the number of SVs discovery and corresponding precision were calculated with caller cuteSV or SVIM after aligner pbmm2 (for deletion) or NGMLR (for insertion) (Fig. 5a-b, d-e, Additional file 2: Tables S8). The count of SVs, both deletion and insertion, increased rapidly with increasing depth and gradually tended to be saturated (Fig. 5a, d). Unexpectedly, the precision had almost no variability with the sequencing depth increasing to ~6.6X (Fig. 5b, e). The extremely slight decrease of the precision had confirmed that using deeper HiFi data could lead to many false positives in keeping with PacBio CLR and Oxford Nanopore long-reads^19^. Also, it might be caused by the more large-scale SV (>10kb), over the length of a long-read, which were obtained with the sequencing depth increasing (Additional file 1: Figure S11-12). These results demonstrated convincingly that deep sequencing can increase recall clearly, but cannot improve precision effectively.

**Fig. 5.**
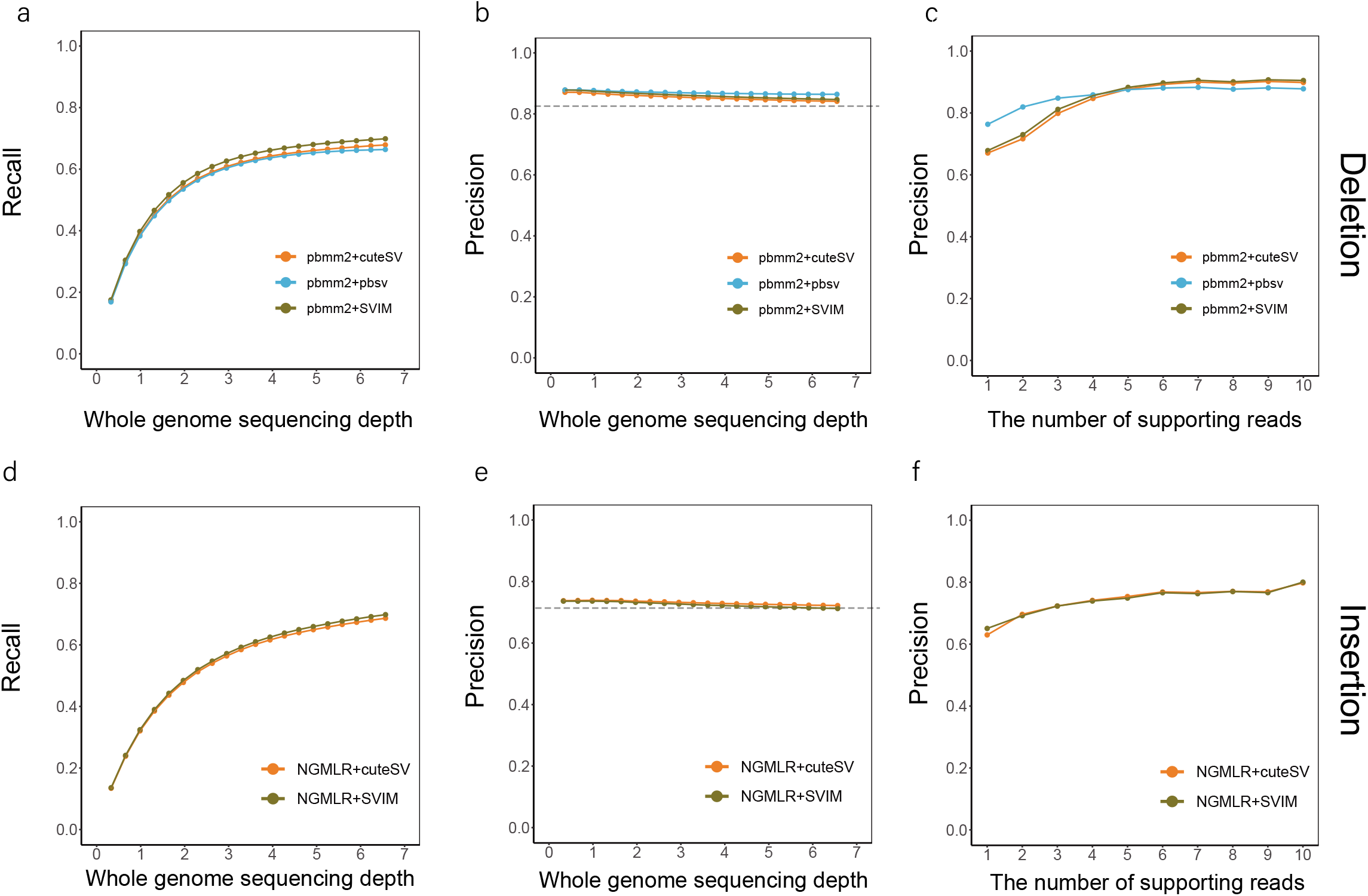
The impact of the sequencing depth and supporting reads variation on SV detection. Precision (**a and d**) and recall (**b and e**) showed the influence of the genome depth after down-sampling from 0.33X to 6.60X. **c and f** The effect of supporting reads on SV accuracy. Caller pbsv after pbmm2 (for deletion) or NGMLR (for insertion) alignment was tagged with blue, caller cuteSV/SIM with red/brown.

### Parameter optimization

The minimum number of supporting reads for a candidate variation is a crucial parameter to call or filter SVs. A recent study revealed that at least 25 long-reads are required to achieve > 80% precision, suggesting that high coverage is essential for SV calling using Oxford Nanopore data^17^. Based on the highly accurate sequence technology, HiFi reads may obtain more reliable results using less supporting long-reads in theory. Hence, we further evaluated the influence of the number of supporting reads on calling accuracy.

The precision values varied from 0.67 to 0.90 on deletion and 0.58 to 0.82 on insertion with the supporting reads increasing to 10 (Fig. 5c, f, Additional file 2: Tables S9). Significantly, only one long-read that supports a candidate SV was required to achieve > 60% precision. In order to achieve relatively higher accuracy, minimal support reads = 3 should be recommended for parameter setting with the precision of deletion (> 80%) and insertion (> 70%), revealing the remarkable ability to detect SVs using low-coverage HiFi data.

## Discussion

Advances in sequence technology—including, but not limited to, the over 10 kilobases (kb) length and 99.8% accuracy of long HiFi reads—greatly speed up the process of large-scale variation study^18^. Correspondingly, new long-reads aligners and SV callers are springing out constantly. Although recent researches have showed the strong point and shortcoming of the current tools for less-accurate ONT long-reads in the human genome^16,17^, there is no knowledge of comprehensively evaluation for the performance of SV mainstream algorithms based on highly accurate long HiFi reads, especially for the large allopolyploid genome in the plant.

In this study, we performed re-sequencing of multiple ploidy genomes using the PacBio CCS method and designed a generally applicable workflow for comparing the precision and recall of single or combining SV sets against the base-level SV truth set utilizing the data integration of multiple sequence technologies. Given the significant differences in aligner selection on SV calling, F-test showed that aligners could explain the higher proportions of total variance compared to callers, suggesting that the performance of SV detection varies depending on the long-read aligners rather than the callers, particularly in deletion identification and SV accuracy (Fig. 3, Tables 1). It also means that a more effective aligner is urgently needed to be developed for getting accurate and comprehensive SV data.

Based on evaluation results, we found that caller cuteSV or SVIM should be recommended as benchmark callers, unrelated to SV type or ploidy level (Fig. 4, Additional file 1: Figure S4, S5-10). However, the selection of aligner obviously differs for SV type with pbmm2 or NGMLR for deletion and insertion detection, respectively (Fig. 4, Additional file 1: Figure S4-5). Besides, high confidence result of insertion could be obtained by the intersection of SV sets, but not in deletion, at the cost of a decline in the number of insertions (Fig. 4, Additional file 1: Figure S6). And the union of SV sets could dramatically improve recall values, both deletion and insertion, while its precision share declined slightly (Fig. 4, Additional file 1: Figure S7). In particular, the more detailed recommendation for users is listed in Table 2.

**Table 2.** Benchmark tools for SV detection in the allopolyploid genome.

| Ploidy | PAV type | Result | Benchmark tools |  | Performance |  |
| --- | --- | --- | --- | --- | --- | --- |
|  |  |  | aligner | caller | precision | recall |
| AABBDD<br>(2n = 6X = 42) | Deletion | Single caller | pbmm2 | cuteSV / SVIM / pbsv | 0.92/0.92/0.93 | 0.47/0.48/0.46 |
|  |  | High-confidence |  | cuteSV & SVIM & pbsv | 0.93 | 0.44 |
|  |  | High-sensitivity |  | cuteSV U SVIM U pbsv | 0.92 | 0.48 |
|  | Insertion | Single caller | NGMLR | cuteSV / SVIM | 0.87/0.87 | 0.75/0.76 |
|  |  | High-confidence |  | cuteSV & SVIM | 0.87 | 0.74 |
|  |  | High-sensitivity |  | cuteSV U SVIM | 0.87 | 0.77 |
| AABB<br>(2n = 4X = 28) | Deletion | Single caller | pbmm2 | cuteSV / SVIM / pbsv | 0.85/0.85/0.86 | 0.49/0.50/0.48 |
|  |  | High-confidence |  | cuteSV & SVIM & pbsv | 0.87 | 0.47 |
|  |  | High-sensitivity |  | cuteSV U SVIM U pbsv | 0.85 | 0.51 |
|  | Insertion | Single caller | NGMLR | cuteSV / SVIM | 0.80/0.79 | 0.70/0.71 |
|  |  | High-confidence |  | cuteSV & SVIM | 0.80 | 0.68 |
|  |  | High-sensitivity |  | cuteSV U SVIM | 0.80 | 0.72 |
| DD<br>(2n = 2X = 14) | Deletion | Single caller | pbmm2 | cuteSV / SVIM / pbsv | 0.84/0.85/0.86 | 0.68/0.70/0.67 |
|  |  | High-confidence |  | cuteSV & SVIM & pbsv | 0.88 | 0.64 |
|  |  | High-sensitivity |  | cuteSV U SVIM U pbsv | 0.84 | 0.70 |
|  | Insertion | Single caller | NGMLR | cuteSV / SVIM | 0.72/0.71 | 0.69/0.70 |
|  |  | High-confidence |  | cuteSV & SVIM | 0.72 | 0.67 |
|  |  | High-sensitivity |  | cuteSV U SVIM | 0.72 | 0.71 |

Another key issue that has to be considered is how to maximize low-coverage data mining under insuring SV accuracy without adding research cost. De Coster et al. reported that either at least 25 long-reads or ~ 8X genome coverage was required to achieve > 80% precision, meaning that the high coverage is essential for SV calling by Oxford Nanopore PromethION sequencing^17^. It is so rejuvenating that the PacBio CCS method can obtain the same–or better–results from only 3 long HiFi reads than ONT data. Even minimal support read = 1 also enables SVs to achieve the precision of deletion (~ 76%) and insertion (~ 65%), revealing the outstanding ability of detecting SVs using low-coverage HiFi data (Fig. 5c, f). In addition, the increase of genome coverage does not appear to affect improvement in precision from 0.33X to 6.60X, which provides strong evidence that deep sequencing can increase recall clearly, but cannot improve precision effectively (Fig. 5b, e). Aiming at the research demand, not the big data, we anticipate that these findings will be used widely to accelerate genomics studies of the PacBio CCS method at the population level.

## Conclusion

No previous study has comprehensively evaluated the performance of the major SV aligner and caller using the PacBio high fidelity (HiFi) reads. This study provided a schematic workflow with wide availability for evaluating the SV detection algorithms in terms of precision and recall. Our results revealed that the performance of SV detection varied depending on the long-read aligners rather than the SV callers. Caller cuteSV or SVIM after pbmm2 (for deletion) or NGMLR (for insertion) alignment should be recommended as benchmarking SV software, unrelated ploidy level. Furthermore, we characterized the impact on the performance of genome coverage and parameter setting for low-coverage data mining. Predictably, this study will facilitate widespread applications of PacBio HiFi sequencing technology for population-scale studies.

## Methods

### Sample preparation and PacBio circular consensus sequencing (CCS)

To facilitate the study of genome-wide identification of SVs in different wheat accessions, we collected three ploidy levels (AABBDD, AABB from *Triticum*; DD genome from *Aegilops*). All three samples were planted in growth chambers. The tender leaves were divided into two equal parts, one half for next-generation sequencing (NGS) on Illumina NovaSeq 6000 system and the other half for Circular Consensus Sequencing (CCS) on PacBio Sequel II system.

### Data processing

#### Reads alignment

By applying our previously designed pipeline for cross-ploidy genetic variation discovery, the NGS/CCS data were mapped to the corresponding wheat reference genome (IWGSC RefSeq v1.0) using short-reads aligner (BWA-MEM) and long-reads aligner (pbmm2, NGMLR^19^, or Minimap2^20^) with default parameters, respectively. The bam files were filtered (unique mapping with mapping quality ≥ 20) and sorted using samtools (version 1.9).

#### SV calling pipeline

SV calling, using pbsv (version 2.3.0), cuteSV (version 1.0.9) ^21^, SVIM (version 1.4.2)^22^, and Sniffles (version 1.0.11) ^19^, was performed following the recommended parameters with minor modifications. For most SV callers, the minimum number of reads was setting 10 as the default. However, the highly accurate HiFi reads may obtain more reliable results using less supporting long-reads in theory. On this basis, minimal support read = 1 was set for SV calling.

#### Candidate SV sets filter

SVs for 11 candidate SV sets presenting the following conditions were retained: (1) SV length□≥□50□bp; (2) minimal support long-read ≥ 1; (3) SVs passing the quality filters suggested by callers (flag PASS).

### The base-level SV truth set construction

#### Bin-deletion method for deletion true set

For every given deletion from the above 11 candidate SV sets, read depth was first calculated by NGS data with mosdepth (version 0.2.6) and the discordant alignment features were collected using samtools (FLAG 1294) and the script “extractSplitReads_BwaMem” developed by lumpy-sv. Due to large amounts of “N” in the wheat reference genome, we further calculated the “adjustDepth” as follows:

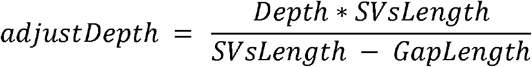

We further investigated the distribution of “adjustDepth”, following Poisson distribution. To obtain the more accurate true set, we selected “adjustDepth = 0” as golden standard. Combining with the discordant alignment features, every given deletion was evaluated to determine if they are true. All true deletions, a higher resolution for breakpoints, obtained by the above method were merged and formed the base-level deletion truth set.

#### Paragraph genotyping method for insertion true set

Given the success of genotyping tools of large-scale variation, Paragraph, an accurate genotyper for short-read sequencing data, was used to validate the insertion dataset that had been mined by 11 candidate SV sets. Previous work usually used the maximum allowed merging distance of 500 or 1000 bp distance, resulting in a decline in the number of SVs and imprecise breakpoint position. Considering the record difference of position for the same insertion among SV callers, 2-bp distance in the left and right breakpoints was chose as the maximum allowed merging distance. All true insertions, obtained by Paragraph, were merge using SURVIVOR ^29^ and formed the base-level insertion truth set.

### Evaluation of the SV detection accuracy

#### Evaluation of single SV call sets

To evaluate the performance of combinations of the aligners and SV callers, the performance for 11 candidate SV sets was assessed against the base-level SV truth set using surpyvor (version 0.6.0) ^17^, a powerful tool for the calculation of precision-recall-F-measure metrics.

#### Evaluation of combining SV call sets

High-confidence or sensitivity SV call sets could be obtained by intersection or union of multiple SV callers. We first constructed the combining SV call sets and then analyzed the performance for each set, using SURVIVOR and surpyvor, respectively.

#### Statistical analysis for SV detection accuracy

Precision (Pr) and recall (Rc) were calculated as follows:

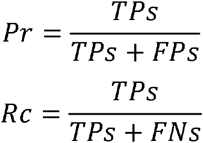

The F-measure (F) is the harmonic mean of precision and recall, which was calculated as follows:

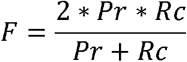

### Analysis of sequencing depth on precision

To study the relationship between the sequence coverage and precision, we randomly down-sampled the sequencing data of the DD sample with 20 gradients from 5% to 100% using Samtools (version 1.8). Following the above method, each coverage sample was evaluated against the base-level SV truth set after reads alignment, SV calling and filter.

## Declarations

### Ethics approval and consent to participate

Not applicable.

### Consent for publication

Not applicable.

### Availability of data and materials

The raw sequence data were deposited in the Genome Sequence Archive (GSA) (https://ngdc.cncb.ac.cn/gsa/) under accession numbers CRA004631.

### Authors’ contributions

FL designed and supervised the research. ZZ, JZ and FL developed the general schematic workflow and performed data analysis. LK, XQ, BN and XF helped with data analysis. AB, XZ, DX, JW and CY collected plant materials. ZZ, JZ and FL wrote the manuscript. All authors discussed the results and commented on the manuscript.

## Acknowledgments

We thank all members (Hua Zhang, Jun Xu, Yafei Guo, Ming Zhang, Xiaohan Yang, Song Xu, Xinyue Song, Jiayu Dong and Liping Jiang) for their valuable comments in Fei Lu’s group, Institute of Genetics and Developmental Biology, Chinese Academy of Sciences.

## Authors’ information

State Key Laboratory of Plant Cell and Chromosome Engineering, Institute of Genetics and Developmental Biology, Innovative Academy of Seed Design, Chinese Academy of Sciences, Beijing, China.

Zhiliang Zhang, Jijin Zhang, Lipeng Kang, Xuebing Qiu, Beirui Niu, Aoyue Bi, Xuebo Zhao, Daxing Xu, Jing Wang, Changbin Yin, Xiangdong Fu, Fei Lu

University of Chinese Academy of Sciences, Beijing, China.

Zhiliang Zhang, Jijin Zhang, Lipeng Kang, Xuebing Qiu, Aoyue Bi, Xuebo Zhao, Daxing Xu, Xiangdong Fu, Fei Lu

CAS-JIC Centre of Excellence for Plant and Microbial Science (CEPAMS), Institute of Genetics and Developmental Biology, Chinese Academy of Sciences, Beijing, China.

Fei Lu

Corresponding author

Correspondence to Fei Lu.

## Publisher’s Note

Springer Nature remains neutral with regard to jurisdictional claims in published maps and institutional affiliations.

