## Additional file 1 for "Genotyping of structural variation using PacBio high-fidelity sequencing"

**Additional File 1: Supplementary figures**

**Genotyping of presence/absence variation using PacBio high-fidelity sequencing**

**This PDF file includes:**

Supplementary figures 1 to 11

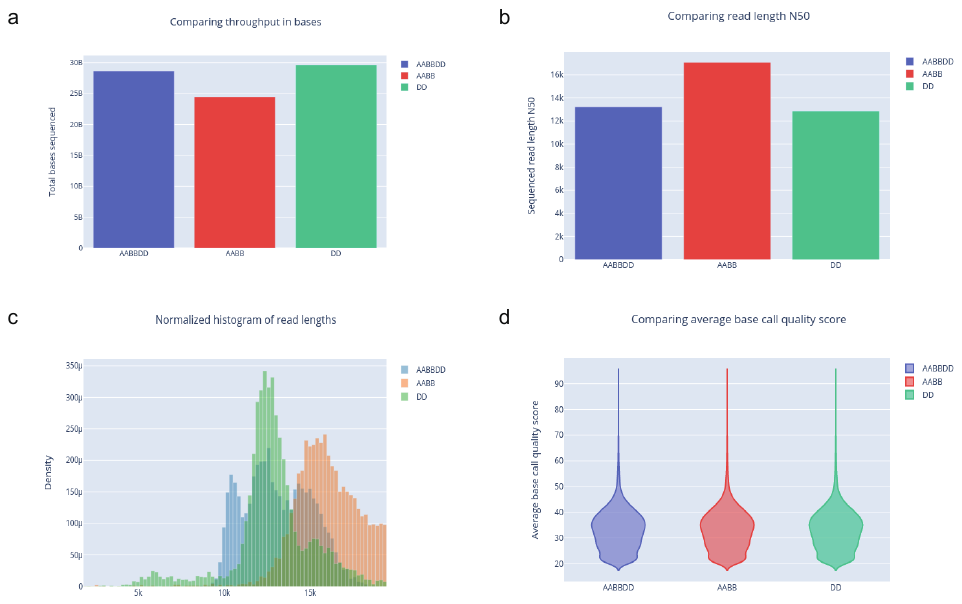

**Figure S1** Summary of all samples in this study. **a - d** showed total output filed (Gb), the length of HiFi reads N50, the distribution of HiFi reads length, and the average base quality score.

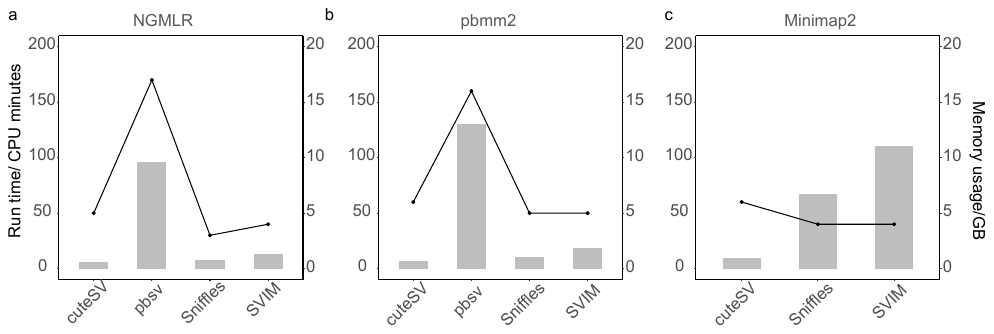

**Figure S2** Run time and memory consumption for PAV detection algorithms. Each of the callers was running using 20 threads.

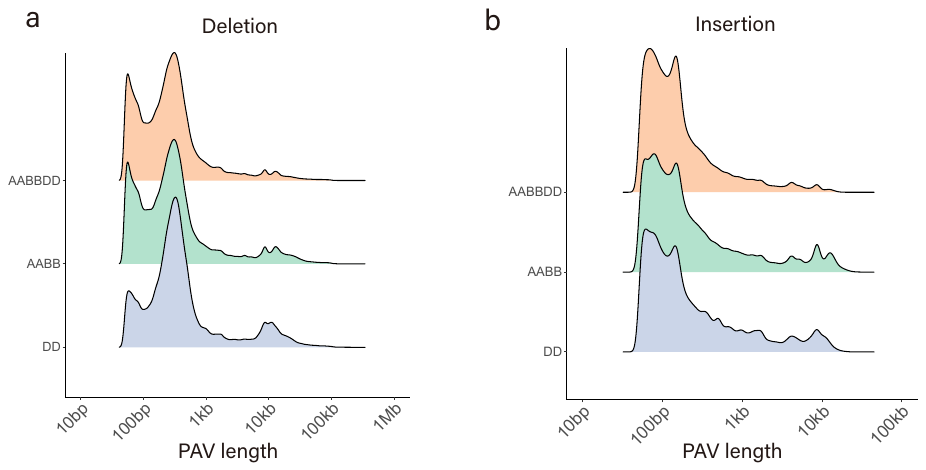

**Figure S3** The length distribution of the base-level PAV truth set. **a and b** for deletion and insertion, respectively.

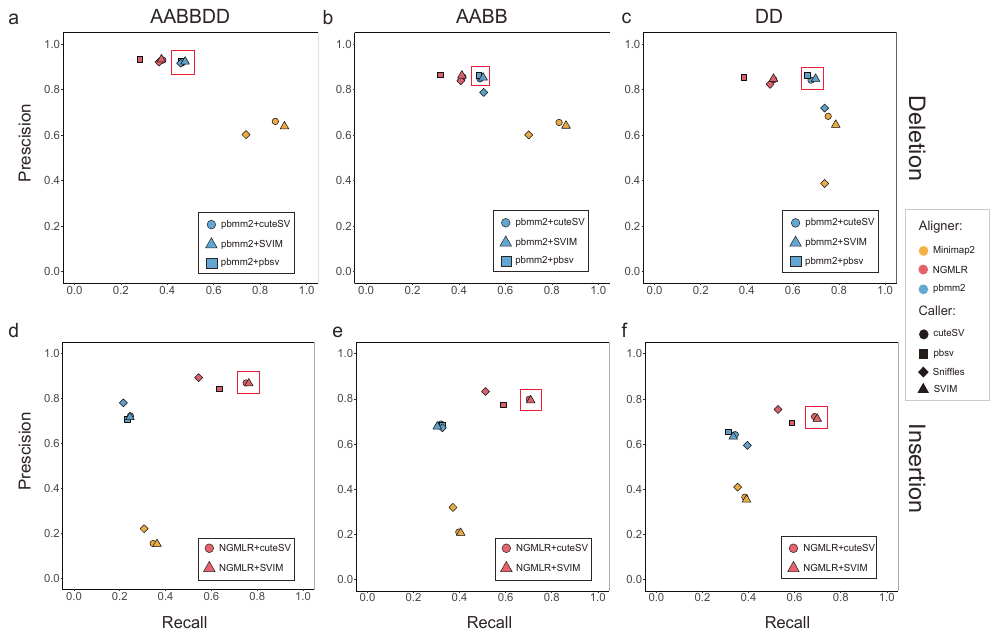

**Figure S4** Precision-recall comparison of single PAV set against the base-level PAV truth set. **a, b and c** indicated precision-recall comparison of deletion in AABBDD, AABB and DD samples. The red box showed the benchmarking tools, cuteSV/SVIM/pbsv after pbmm2 alignment, for deletion. **d, e and f** indicated precision-recall comparison of insertion in AABBDD, AABB and DD samples. The red box showed the benchmarking tools, cuteSV/SVIM after NGMLR alignment, for insertion.

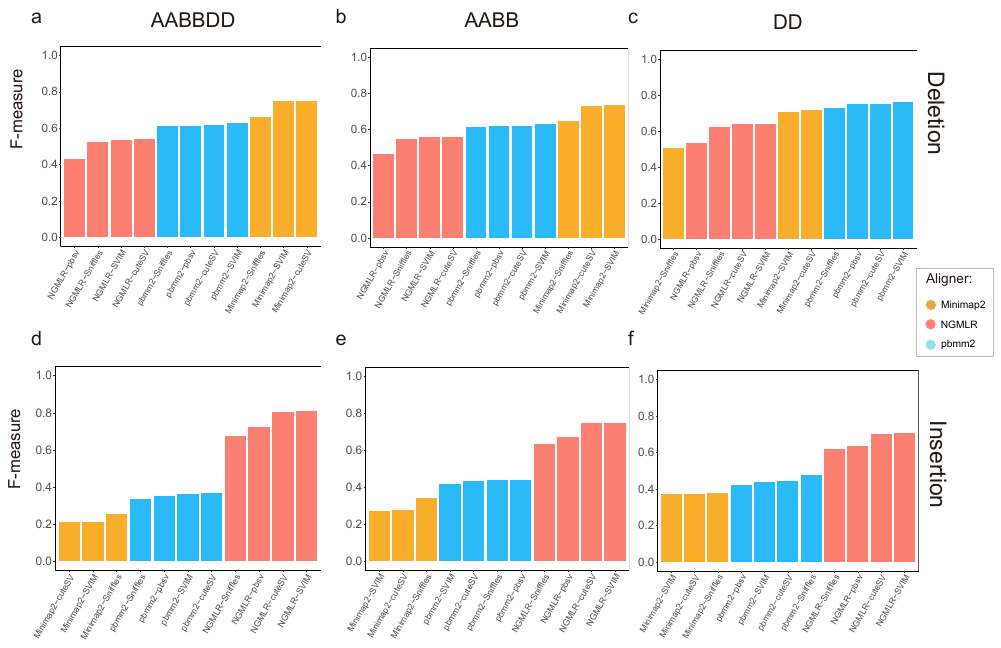

**Figure S5** F-measure comparison of single PAV sets against the base-level PAV truth set. 11 single-caller PAV sets were obtained by the caller and aligner combination of pairs. The inverse red triangle denoted the maximum F-measure, except the PAV sets based on Minimap2. Aligner Minimap2, NGMLR and pbmm2 were tagged with orange, red and blue, respectively.

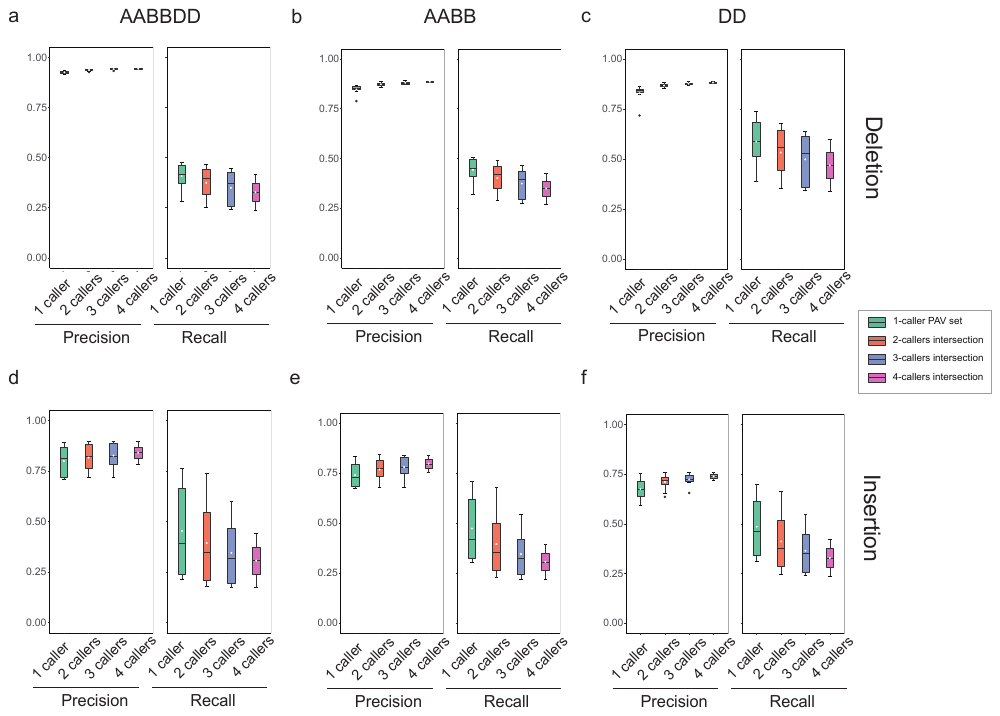

**Figure S6** Precision-recall comparison of high-confidence PAV sets in three ploidy levels**. a - c and d - f** for deletion and insertion, respectively. All PAV sets were obtained by the intersection of different callers based on the same aligner. The white dots represent the mean value.

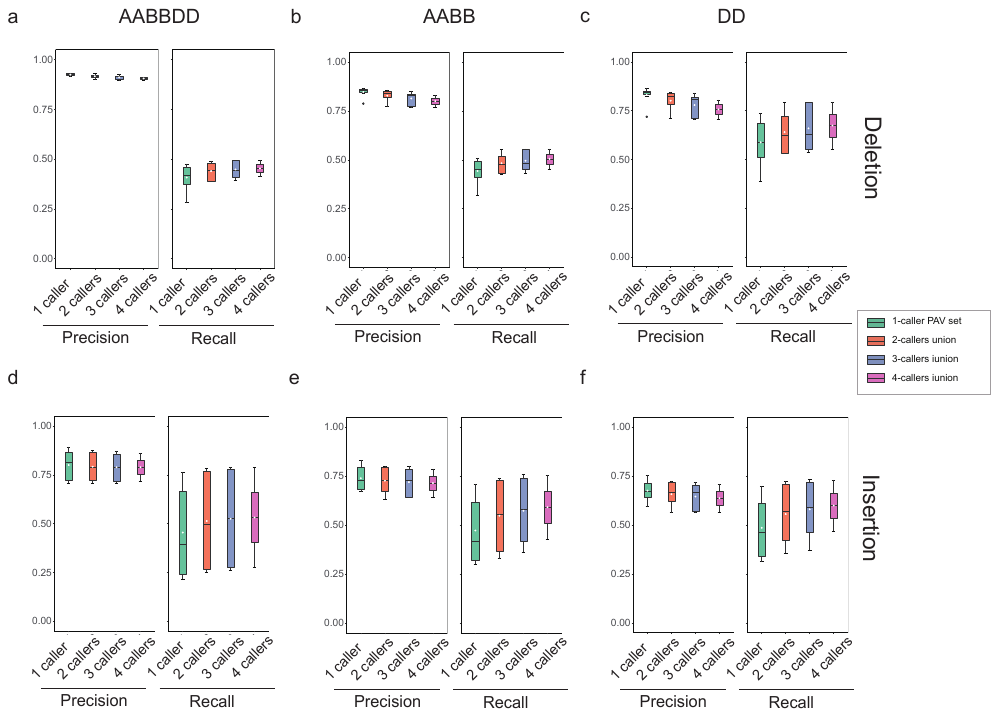

**Figure S7** Precision-recall comparison of high-sensitivity PAV sets in three ploidy levels**. a - c and d - f** for deletion and insertion, respectively. All PAV sets were obtained by the union of different callers based on the same aligner. The white dots represent the mean value.

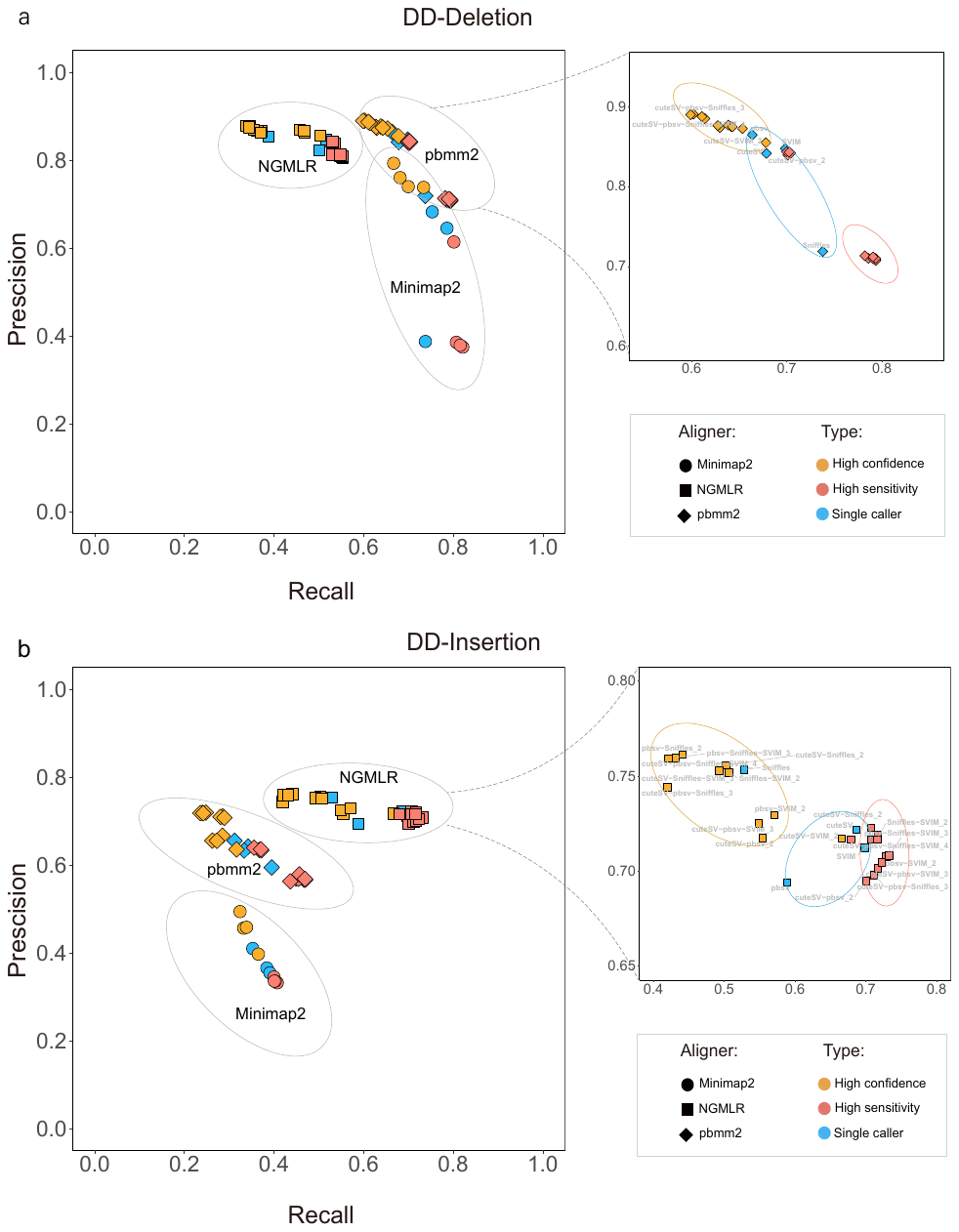

**Figure S8** Comprehensive evaluation of 63 PAV sets in diploid (DD) genome. **a** (deletion) **and b** (insertion) Precision-recall graph of single/combining PAV call sets against the base-level PAV truth set. Aligners are represented by symbols, and multiple set sources are represented by colors as specified in the legend.

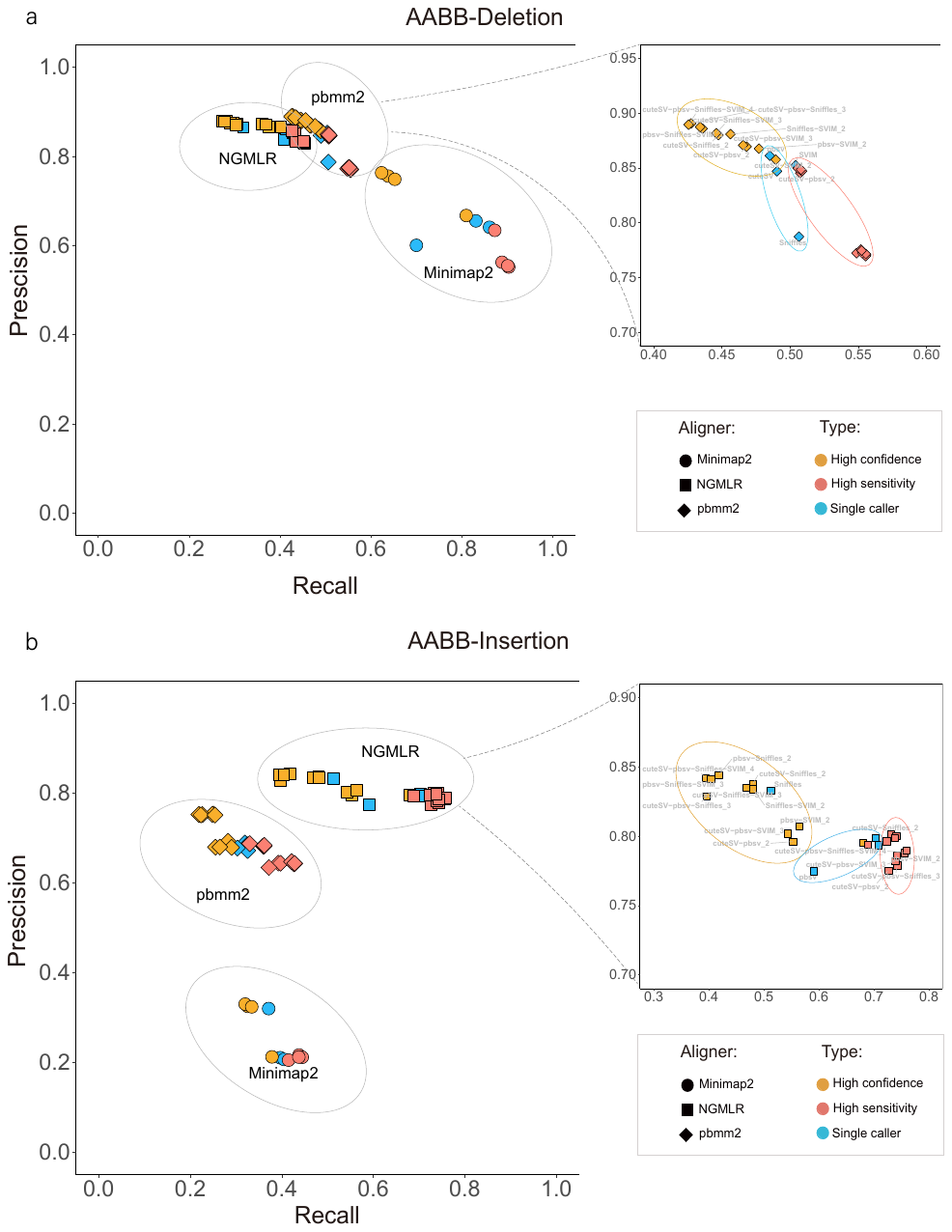

**Figure S9** Comprehensive evaluation of 63 PAV sets in tetraploid (AABB) genome. **a** (deletion) **and b** (insertion) Precision-recall graph of single/combining PAV call sets against the base-level PAV truth set. Aligners are represented by symbols, and multiple set sources are represented by colors as specified in the legend.

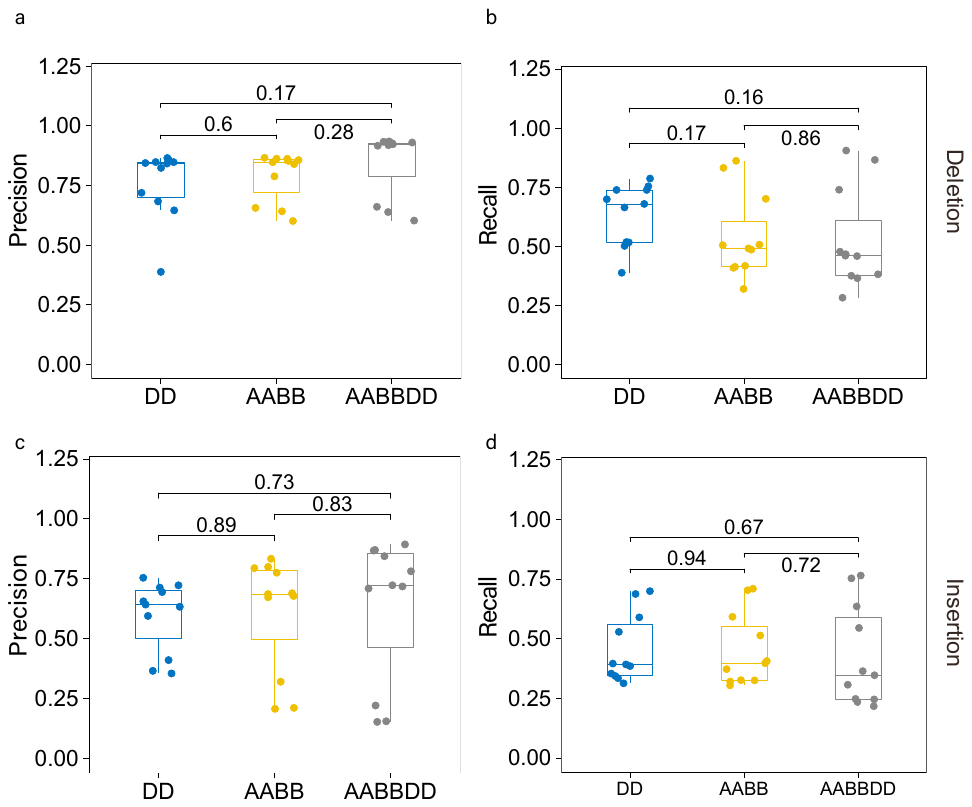

**Figure S10** The performance comparison of single-caller PAV sets in different diploidy levels. **a, b** (deletion) **and c, d** (insertion) There was no significant difference on precision/recall in DD, AABB, AABBDD genome.

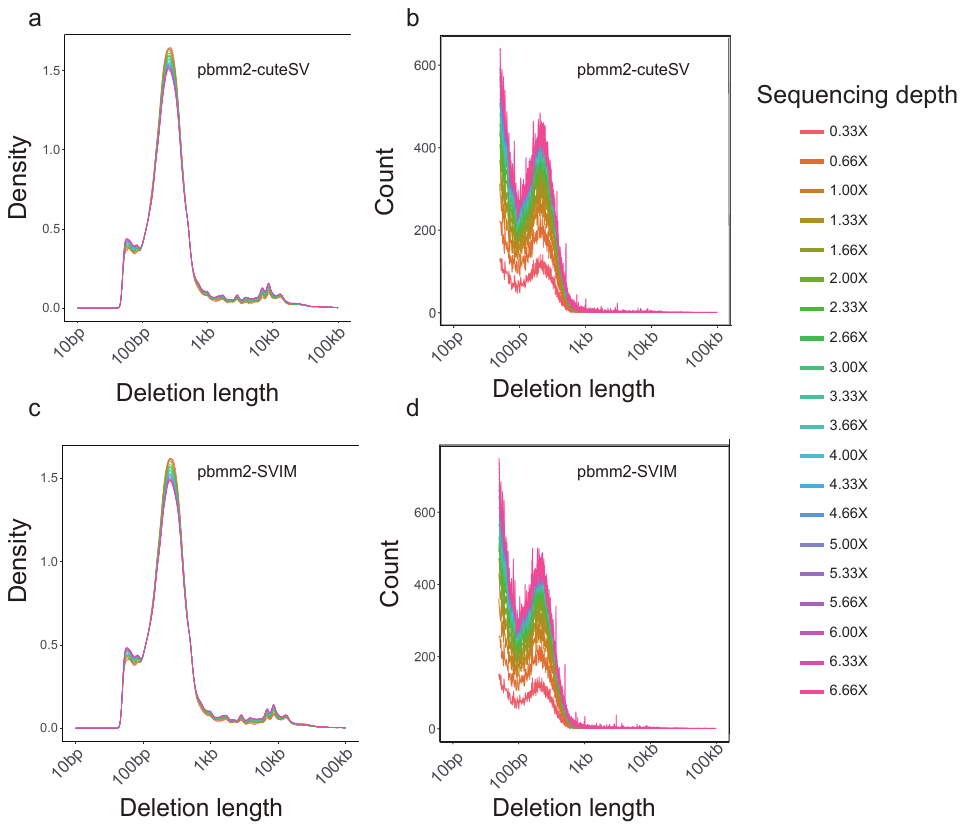

**Figure S11** The impact of the sequencing depth on the length distribution of deletion. The more large-scale deletion (>10kb), either cuteSV or SVIM after pbmm2 alignment, were detected with the sequencing depth increasing.

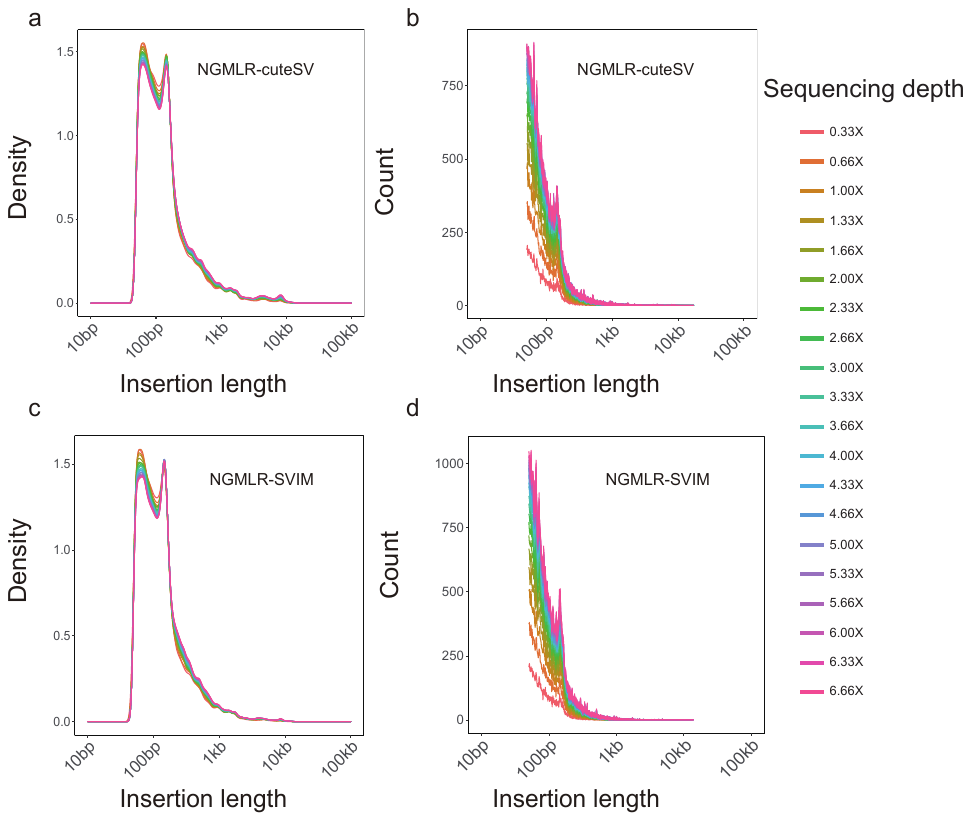

**Figure S12** The impact of the sequencing depth on the length distribution of insertion. The more large-scale insertion (>10kb), either cuteSV or SVIM after NGMLR alignment, were detected with the sequencing depth increasing.
